# The mutation effect reaction norm (Mu-RN) highlights environmentally dependent mutation effects and epistatic interactions

**DOI:** 10.1101/2021.09.23.461533

**Authors:** C. Brandon Ogbunugafor

## Abstract

Since the modern synthesis, the fitness effects of mutations and epistasis have been central yet provocative concepts in evolutionary and population genetics. Studies of how the interactions between parcels of genetic information can change as a function of environmental context have added a layer of complexity to these discussions. Here I introduce the “mutation effect reaction norm” (Mu-RN), a new instrument through which one can analyze the phenotypic consequences of mutations and interactions across environmental contexts. It embodies the fusion of measurements of genetic interactions with the reaction norm, a classic depiction of the performance of genotypes across environments. I demonstrate the utility of the Mu-RN through the signature of a “compensatory ratchet” mutation that undermines reverse evolution of antimicrobial resistance. More broadly, I argue that the mutation effect reaction norm may help us resolve the dynamism and unpredictability of evolution, with implications for theoretical biology, genetic modification technology, and public health.

## I. INTRODUCTION

Modern perspectives in evolutionary genetics are increasingly driven by notions that complex traits are the product of interactions between genes [1, 2]. In many ways, these are renditions of classical debates surrounding the eminence of gene interactions that were a feature of the modern synthesis [3, 4, 5]. At one level, the debates have not changed much during the last century, still defined by a simple question– how many different actors do we need to consider in order to understand the relationshp between genotype and phenotype? Can we understand meaningful changes in phenotypes by studying biology one-mutation-at-a-time? Or do we need better models for how mutations interact with each other, and/or the environment?

The importance of understanding interactions between mutations has emerged as its own area of evolutionary genetics, to the tune of several different (but related) concepts that are studied under the umbrella concept of epistasis. One of these concepts, “physiological epistasis,” has been defined as “any situation in which the genotype at one locus modifies the phenotypic expression of the genotype at another [6].“An expansive literature exists that has examined physiological epistasis in adaptive landscapes [7, 8, 9], with respect to protein biophysics [10, 11, 12], in terms of genomic architecture [13, 14], and many other arenas.

This broader notion that mutations may interact with other parcels of genetic information in a cryptic, spurious fashion casts a shadow over much of modern genetics [15, 16, 17, 18], and may contribute to phenomenon like phantom heritability [19, 20, 21]. Relatively underexplored in conversations about how interactions manifest in complex phenotypes are theoretical treatments of how environmental gradients may influence the interactions between mutations or SNPs.

Conveniently, an abstraction exists in the evolutionary biology and ecology canons—the reaction norm (also known as the “norm of reaction”)—to describe how the environment shapes the performance (phenotype) of genotypes [22]. The reaction norm is widely applied in quantitative genetics [23, 24], in discussions of phenotypic plasticity [25, 26, 27], and other subtopics.

While several studies have examined how environments can tune nonlinear interactions between mutations [28, 29, 30,31, 32, 27, 33], there have been few formal attempts to integrate details of the environment into measurements of mutation effects and interactions. In this study, I introduce the “mutation effect reaction norm,” an abstraction that combines the reaction norm with mutation effects and physiological epistasis. It demonstrates how the strength and nature of interactions can change appreciably across environmental contexts of various kinds. To demonstrate its utility, I explore data sets corresponding to a collection of alleles associated with antimicrobial drug resistance. I analyze these data using the mutation effect reaction norm framework and diagnose the signature of a “compensatory ratchet” mutation whose effect is specific to environment.

Summarizing, I discuss how this abstraction is relevant in many problems where the effects of individual mutations are influenced by environments, including genetic modification, public health and biomedicine. More broadly, I use the concept to emphasize the importance of more detailed biographies of mutation interactions in present and future attempts to capture the shape of molecular evolution.

## II. METHODS

### A. Data Sets

While much of the argument surrounding the utility of the Mu-RN is conceptual, I thought that it would be most effective to demonstrate how it works using real world data and analyses. In this way, the ideas are less abstract, and the reader can observe firsthand how they can be applied.

In this study I decided to study mutation effects and interactions involved in the evolution of antimicrobial resistance. I utilized two data sets, each a combinatorially complete set, where suite of mutations within a locus (a protein in this case) are engineered in all possible combinations. While this data structure is not necessary to measure interactions and epistasic effects, it facilitates the use of certain transparent, established algebraic formulations.

#### A note on the use of “reaction norm.”

I should mention that use of the term “reaction norm” describes depictions of how the alleles in this set perform across drug environments. Very similar analyses and descriptions were used in prior studies [34], but without using the “reaction norm” descriptor.

The traits of interest in this study are growth rates of the alleles in different concentrations of pyrimethamine and cycloguanil, antifolate drugs used to treat malaria [35]. I examined 16 alleles composed of combinations of four mutations (N51I, C59R, S108N, I164L; 2^4^ = 16 alleles)) in *Plasmodium falciparum* (a cause of malaria) dihydrofolate reductase (DHFR, an essential enzyme). All 16 alleles have growth rate values across a gradient of drug concentrations (10^−3^ *μM* to 10^5^ *μM*). These data arise from a set utilized in previous studies that examined the evolution of resistance to antimalarials [36, 34]. Also note that while we use growth rates as our main phenotype, the methods described here can be used to study fitness measurements of various kinds, including relative fitness.

### B. A note on methods to measure interactions

Many methods exist for measuring the strength of interactions between mutations in empirical datasets. Questions surrounding which methods are appropriate are similar to many statistical questions surrounding how to disentangle nonlinear effects in complex systems (biological or other): the shape, scope, and size of the data dictate which analyses are most appropriate. Furthermore, epistasis might be described as an idea whose definitions are at least partly based on how it is measured. For example, some methods consider how noise can conflate the measurement of resistance [37, 38], accommodate non-binary encoding or gaps in data [39], interrogate the limits of regression methods [40, 41], or measure marginal epistasis across large genomic data sets [42]. These methods can be considered more ideal for certain questions or data structures. However, a rigorous treatment of methods used to measure epistasis is beyond the scope of this manuscript.

#### An example method: The Walsh-Hadamard transform

In this study, I used the Walsh-Hadamard transform, which computes a coefficient corresponding to the magnitude and sign of an interaction between mutations with respect to a phenotype. It was pioneered for use in the study of higherorder epistasis in a 2013 study that both provided a primer for the calculation and analyzed several combinatorially complete data sets [7]. It has since been further applied to study of higher-order epistasis across a larger sampling of empirical data sets [43].

The Walsh-Hadamard transform implements phenotypic measurements into a vector, then a Hadamard matrix, which is scaled by an additional diagonal matrix and is used to act on this phenotypic vector. The result is a set of coefficients that measure the degree to which the genotype-phenotype map (perhaps described in the guise of an adaptive landscape) is linear, or second order, third, and so forth. For more rigorous discussions of the method, I encourage readers to engage several published manuscripts—especially Weinreich et. al. (2013) [7] and Poelwijk et al. (2016) [44] — each of which explore the methods and their related issues in greater detail. For clarity, I will describe selected aspects of the method in this manuscript.

As the data examined by the Walsh-Hadamard transform are combinatorially complete, one can represent the presence or absence of a given mutation by a 0 or 1 at a given locus. For example, one can represent a wildtype variant of a gene as 0000. In this one scenario, the mutations at each of four sites (e.g. the four mutations corresponding to antifolate resistance *Plasmodium falciparum* dihydrofolate reductase) [45, 46] are, N51I, C59R, S108N, and I164L. For those unfamiliar with this notation: the number corresponds to the location in the protein, and the letters on each side of the number correspond to single-letter amino acid abbreviations for the variants at that site. For example, N51I corresponds to an asparagine to isoleucine mutation at the 51st amino acid in DHFR). The quadruple mutant, IRNL, would be encoded as 1111 in this scenario.

The full data set consists of a vector of phenotypic values (growth rate in the presence of two different antifolates) for all possible combinations of mutations (for 16 alleles in total):

NCSI, NCSL, NCNI, NCNL, NRSI, NRSL, NRNI, NRNL, ICSI, ICSL, ICNI, ICNL, IRSI, IRSL, IRNI, IRNL

These can be depicted in binary notation as the following:

0000, 0001, 0010, 0011, 0100, 0101, 0110, 0111, 1000, 1001, 1010, 1011, 1100, 1101, 1110, 1111

This vector of phenotypes, denoted by *p* (arranged *numerically* as in the order presented above), is multiplied by a (16 × 16) square matrix, which is the product of a diagonal matrix *V* and a Hadamard matrix *H*. These are defined recursively:

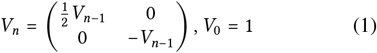

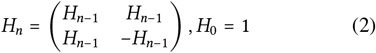

n is the number of loci (n = 4 in this *Plasmodium falciparum* DHFR setting corresponding to resistance mutations). For *n* = 4, these matrices can be depicted as follows:

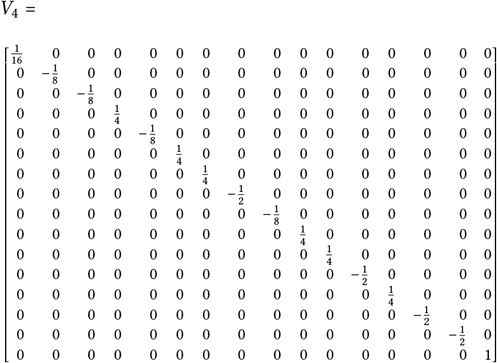

and

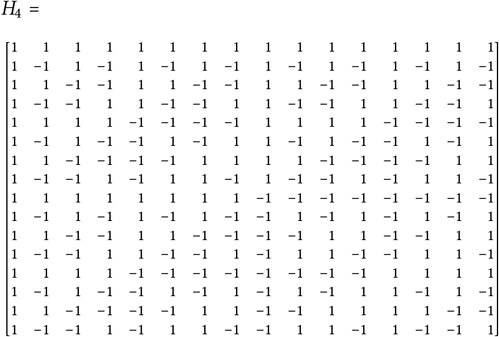

These are combined with the phenotype vector as follows:

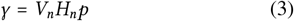

Where *V*_*n*_ and *H*_*n*_ are the matrices described in equations 1 and 2 above and *γ* is the Walsh coefficient vector, the measure of the interaction between mutations. Using this formulation, I compute *γ* values for every possible interaction between bits in each string.

In addition to the aforementioned references where this approach was introduced and described in good detail [7, 44], this method is also made available for exploration in the Supporting Information. It contains a spreadsheet that outlines the calculation and provides a means for inexperienced users to calculate interaction coefficients. While this is not a substitute for learning the methods from their proper sources, it does provide a simple way for those interested to perform these calculations on data of a certain structure.

Having outlined the method used to calculate the Walsh-Hadamard coefficient, I must be clear about the interpretation. The Walsh coefficient corresponds to the average effect of a given mutation effect (first order, pairwise, etc.) across all cognate genetic backgrounds. Negative values for an effect suggest that the average effect is adverse for a given phenotype in a given setting, positive if it has a beneficial effect on a phenotype (e.g., antibiotic resistance). Please note that the idea of “adverse” or “beneficial” in this context is simply a description of its quantitative impact, and not a biological (or ethical/moral) interpretation of the “goodness” of an effect.

One limitation of the iteration of the Walsh-Hadamard transform used in this study is that it requires combinatorial data sets, where an often small set of mutations are constructed in all possible combinations. Another limitation is that it can only accommodate two variants (amino acid substitutions in this case) per locus. For example, if one wanted to measure the higher-order interactions between 4 mutations within a gene, one would need 2^*L*^ = 16 individual measurements, with L corresponding to the number of different mutations whose effects one is interested in disentangling (4 in this case). Another established limitation of the method is that it doesn’t formally incorporate experimental noise. Consequently, its resultant measurement is more consistent with an average of the effect of a mutation or mutation-interactions. Though these limitations reveal that the Walsh-Hadamard transform might be specific to certain datasets, it still applies to many real-world settings, and provides relevant biological insight.

### C. Calculations of higher-order epistasis

Previous studies have examined how higher-order epistasis manifests across empirical adaptive landscapes [7, 43, 30]. “Order” corresponds to the number of actors involved in an interaction. “First-order” would correspond to the effect of single mutations, second-order or “pairwise” interactions would apply to pairs of mutations, and so forth. One can calculate higher-order epistasis using several minor modifications to the Walsh-Hadamard transform method outlined above, very similar to how prior studies carried this out [7].

For example, in a combinatorially complete data set comprising 16 alleles, one can also depict the interactions between individual loci and genetic background using a binary representation (just as one can with whole alleles). In this case, each 0 or 1 represents a locus interaction. To emphasize the distinction in using binary notation both for phenotype and for epistasis coefficients, one should consider using language like *γ*_0000_ for clarity:

*γ*_0000_: zeroth order interaction

*γ*_0001_, *γ*_0010_, *γ*_0100_ & *γ*_1000_ are first-order interactions. For example, this translates to the average effect of the N51I mutation across all possible genetic backgrounds (composed of combinations of the other three loci), those between the C59R mutation and all possible genetic backgrounds, between S108N mutation and all possible genetic backgrounds, and between I164L and the other loci.

Relatedly: *γ*_0011_, *γ*_0101_, *γ*_0110_, *γ*_1001_, *γ*_1100_ & *γ*_1010_ are secondorder or pairwise interactions; *γ*_0111_, *γ*_1011_, *γ*_1101_ & *γ*_1110_ are third-order interactions; and *γ*_1111_ is a fourth-order interaction, the interaction between the four mutations that constitute the quadruple mutant, IRNL. For even more clarity, one can replace the 0s with asterisks (*) to emphasize that binary sites represent mutation interaction effects across all possible genetic backgrounds. For example, the pairwise effect coefficient corresponding to “0110” truly means the average effect of the C59R and S018N mutations, across all other genetic backgrounds. One can depict this effect as “*11*”.

Though the data used in this study are not normalized, it is often prudent to take the absolute value of coefficients, then compute a normalized version of the epistatic coefficients. The normalization standardizes the value so that the analyses might be compared to other data sets. For a given epistatic coefficient *γ*, I define the normalized epistatic coefficient E, as in prior studies of in silico adaptive landscapes [47]:

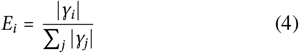

Where the sum over *j* runs over all epistatic coefficients comprised in *γ*. In this study, I only use the absolute values of the epistatic coefficients for all analyses, without normalization. One can average the interaction coefficients within an order to facilitate comparisons between orders (e.g., are third order effects stronger than pairwise effects across environments?). I label these order-averaged effects with the term “absolute mean.” This provides mean values for each order, which calculates the overall contribution of, for example, 1st order effects and higher-order (3^*rd*^ order, 4^*th*^ order, etc.) effects. And one can examine how the order of effects changes across environmental gradients, representing a kind of mutation effect reaction norm for higher-order epistasis.

## III. RESULTS

First, we discuss the reaction norm, the performance (growth rate) of the 16 alleles across drug environments. Second, we will discuss what happens when these alleles are deconstructed into their mutation effects using the procedures outlined in the Method and depicted across environments. This is the anatomy of the Mu-RN, and the most critical aspect of the results. Lastly, we provide a real-world example of a type of problem that the Mu-RN might be applied to: the diagnosis of mutations that have certain effects in specific environments, and serve as “compensatory ratchets” against reverse evolution.

### A. The reaction norm demonstrates the growth rate of alleles across drug environments

Figure 1 depicts reaction norms for a combinatorial set of 16 alleles. The data demonstrate growth rates as a function of concentrations of two different drugs: pyrimethamine (1A, B) and cycloguanil (1C, D).

**Figure 1:**
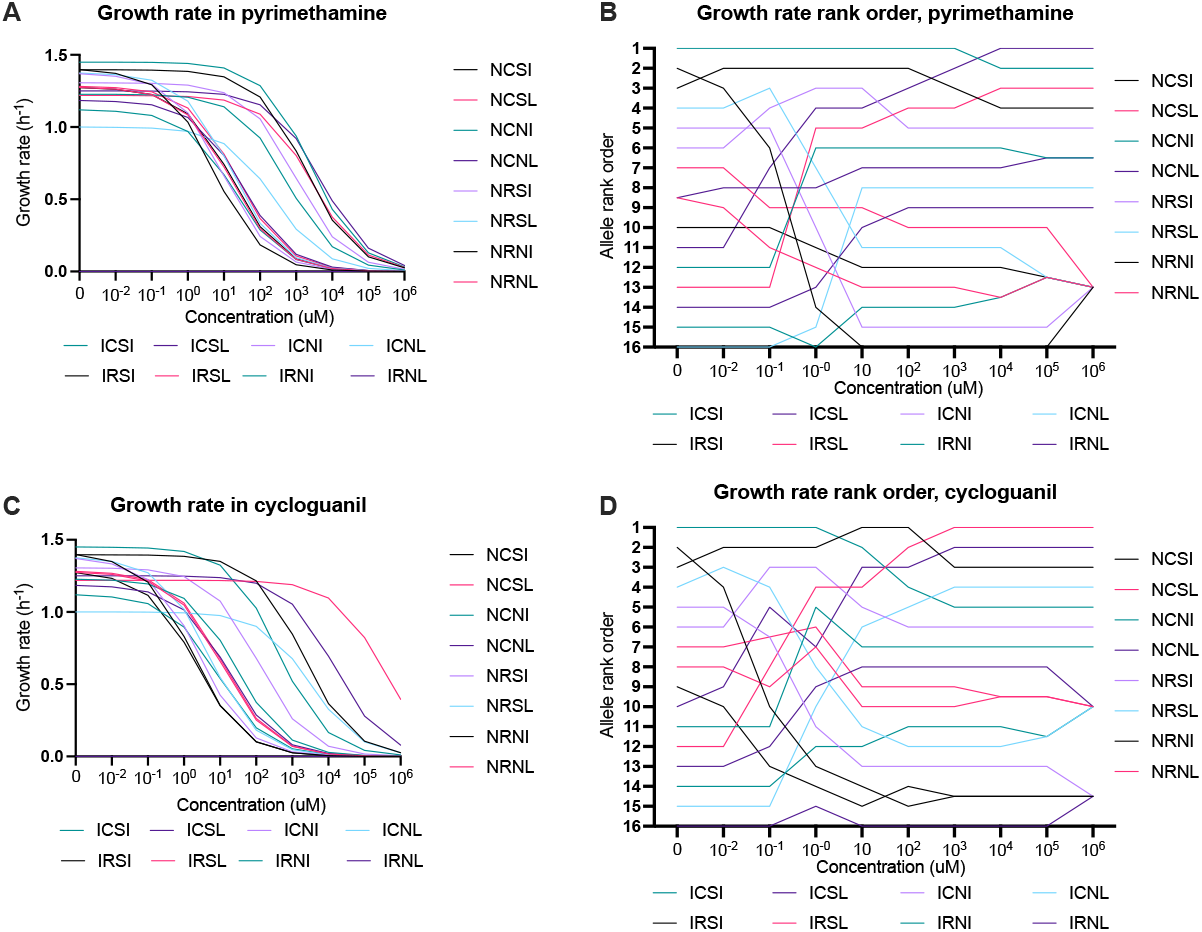
Reaction norms and rank orders. The reaction norm for growth rates corresponding to *Plasmodium falciparum* carrying 16 different alleles of dihydrofolate reductase associated with resistance to antifolates, across drug environments. (A) Reaction norm for growth rate of alleles across drug concentrations of pyrimethamine. (B) Rank order of alleles across drug concentrations in pyrimethamine. (C) Reaction norm for growth rate of alleles across drug concentrations of cycloguanil, and (D) Rank order of alleles across drug concentrations in cycloguanil.

The dynamism of the reaction norms is further encapsulated by depictions of the respective rank orders of alleles in the presence of the two antimalarial drugs, pyrimethamine and cycloguanil (Fig. 1B, D). That the rank order of alleles changes rapidly at some concentrations is a signature of epistasis present in the system, as rank order reflects nonlinear interactions between the mutations that compose the allele [48]. Specifically, note the rapid rank order changes occurring at certain drug concentrations (roughly from 10^−2^*μM* to 10^2^ *μM*, on both pyrimethamine and cycloguanil). The data that compose these reaction norms were previously examined with respect to how drug environments create different evolutionary dynamics (across drug type and concentration) [36, 34].

### B. Exploring how the phenotypic effects of mutations changes according to environment

Mutation effect reaction norms display interactions between mutations for two similar antifolate drugs (pyrimethamine and cycloguanil) along continuous environmental dimensions (Fig. 2A and 2C). Using this approach, we can observe how mutation effects are tuned differently across environments that are at least somewhat similar. That is, pyrimethamine and cycloguanil are both antifolate drugs used to treat malaria infections and have slightly different patterns of mutation effects across contexts, all coefficients converging towards [0] as drug concentrations increase. That is, at a high enough drug concentration, no alleles grow (see the reaction norm results, Figure 1), and so the mutation interactions that compose the alleles in the reaction norm have small effects at the highest drug concentrations.

**Figure 2:**
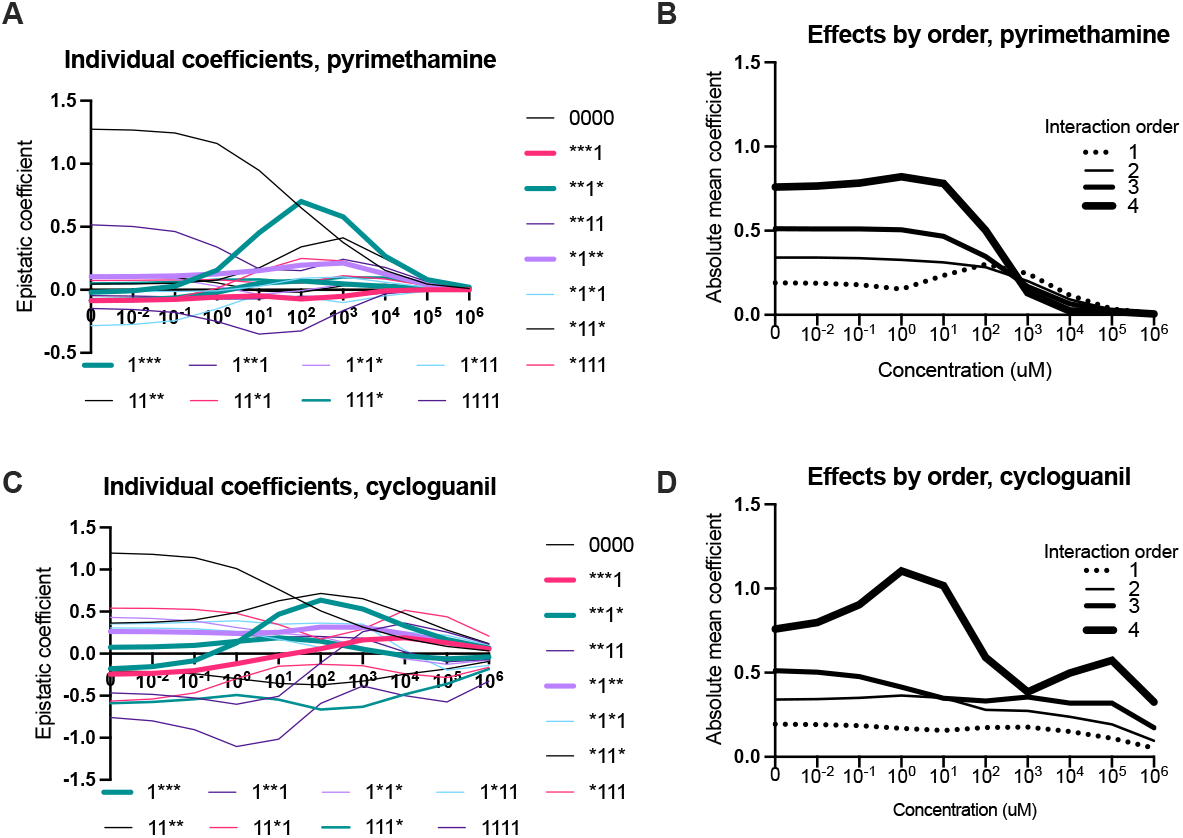
Mutation effect reaction norms (Mu-RN). The mutation effect reaction norm corresponding to the strength of in- teractions. (A) The Mu-RN depicts interactions between individual loci associated with resistance within the *Plasmodium falciparum* dihydrofolate reductase across a breadth of concentrations of pyrimethamine (x-axis title removed for clarity). The binary notation corresponds to interactions between one of four individual loci within the *P. falciparum* dihydrofolate reductase. (B) Mu-RN corresponding to a transformation of the data in (A), whereby the absolute mean of values of all the effects of a certain order are combined, which provides a perspective on how higher-order epistasis varies across environmental context (see Methods for details). (C) A Mu-RN for individual loci interactions effects on growth rate across cycloguanil concentrations (x-axis title removed for clarity). (D) A Mu-RN depiction of higher-order epistasis for resistance across a set of cycloguanil environments. Note that both A and C, the single mutation effects corresponding to **1* is emphasized with a thicker line. This effect, corresponding to the average effect of the S108N mutation across genetic backgrounds and drug environments, is of special interest, as discussed in the main text.

The differences between pyrimethamine and cycloguanil manifest in the topographies of their respective adaptive landscapes[34, 36] and in the shape of their mutation effect reaction norms. For clarity, the average effects of single mutations are emphasized in the Figures 2A and 2C (thicker lines), as they are the coefficients whose interpretation are the most intuitive: the single mutation effect lines provide an average description of how impactful each of the four individual loci are across the drug environments. The values of the remainder of the interactions arise from the formula outlined in the Walsh-Hadamard transform calculation (see equations 1-4) and are more challenging to summarize verbally (see Methods).

To offer a clearer understanding of how environments shape higher-order interactions, I provide a mutation effect reaction norm corresponding to the absolute mean values of mutation effects, organized by order (Fig. 2B and 2D). These represent magnitude differences between orders of effects and communicate the overall presence of higher-order interactions across environmental gradients (the environments being drug type and concentration).

### C. The mutation effect reaction norm highlights the specific signature of “compensatory ratchet” mutations

Of particular interest are the effects of S108N in *P. falciparum* dihydrofolate reductase (Fig. 2A, 2C). The effects of an orthologous mutation were described in a study of reverse evolution of antifolate resistance in *Plasmodium vivax* [49]. Note that in both pyrimethamine and cycloguanil, the mutation effect has a similar pattern: a negative effect at low drug concentrations, with a sign change (from negative to positive) as drug concentrations increase towards 1.0 *μM*. As drug concentrations get very high, mutation effects are low.

This change of of the sign of a mutation effect (from negative to positive) is a signature of a mutation that could be described as “compensatory.” For example, the mutation corresponding to S108N is conditionally beneficial, conferring positive epistatic interactions in high drug concentration environments (Fig. 2A). These mutations restore growth in genetic backgrounds where alleles are growing poorly (generally true in high drug concentrations). This compensatory S108N mutation also serves as a ratchet that undermines the reversal of evolution (from the IRNL quadruple mutant towards the NCSI wildtype in this setting).

Figure 3 further describes how mutation interactions involving the S018N mutation can influence evolution. In 3A, we observe that alleles that contain S018N have a significantly higher growth rate, across all drug concentrations of both pyrimethamine (Kruskal-Wallis: 5A, pyrimethamine, p = 0.0002). As mentioned, the S108N mutation is an ortholog of a mutation, S117N, that has been described as a “pivot” mutation, that both dictates the direction of adaptive evolution, and precludes reversal [49].

**Figure 3:**
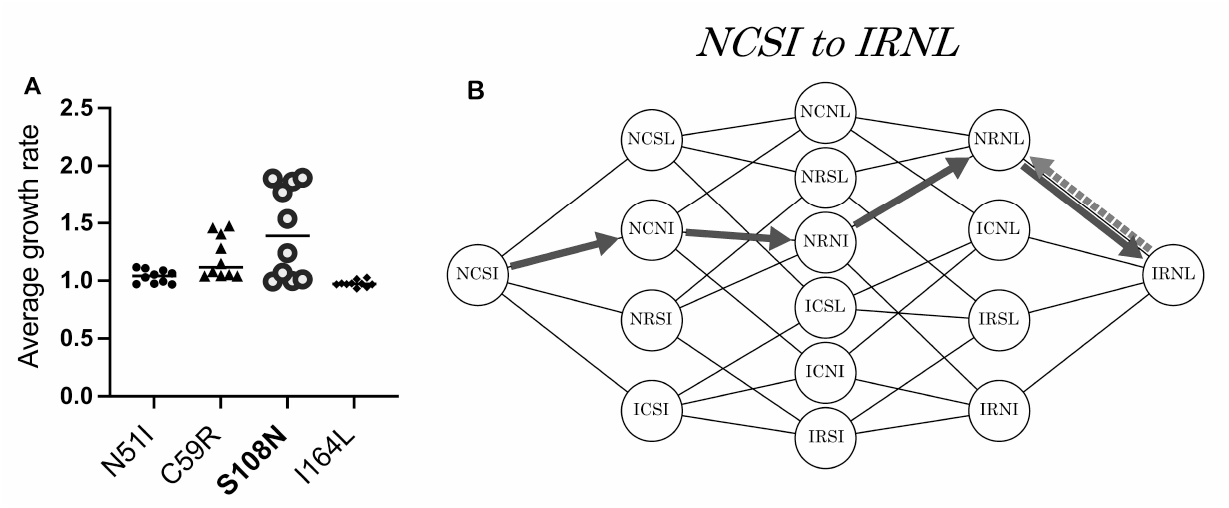
An example application of the Mu-RN: The signature of a “compensatory ratchet” mutation. Here we depict how the S108N mutation’s effect across environments plays a critical role in “forward” evolution and undermines “reverse” evolution. (A) Averaged across pyrimethamine concentrations, alleles containing the S018N mutation grow significantly better than any set of alleles containing any other single mutation (Kruskal-Wallis: 5A, pyrimethamine, p = 0.0002). This is because of the compensatory effects of the mutation at high drug concentrations. (B) The hypercube represents the combinatorial set of 16 alleles as described in the Methods. Based on the rank-order in Figure 1, the predicted pathways of stepwise evolution from the wild-type genotype (NCSI) through the adaptive landscape at a high drug concentration (black arrows) at 10^6^ μ*M*, and reversal in a drugless environment (dashed arrow). In this scenario, the compensatory nature of the S108N mutation provides a “compensatory ratchet,” that helps evolution evolve towards a fitness peak in the high drug concentration (black arrows) but prevents it from reversing towards wildtype (NCSI) in a drugless environment (dashed arrow).

Figure 3B is a hypergraph summary of the predicted evolutionary trajectories in pyrimethamine [34]. Predictions can be made from the rank orders of alleles outlined in Figure 1. That is, starting from NCSI, evolution may follow a path of increasing growth rate, a proxy for reproductive fitness in this setting. Figure 3B depicts “forward” evolution starting from the wild type (NCSI) allele evolving at 10^6^ *μM* pyrimethamine, as summarized in previous studies [34]. In addition, Figure 3B shows the predicted “reverse” evolution trajectory, when the IRNL quadruple mutant evolves in a drugless environment. The preferred trajectories for both forward and reverse evolution in pyrimethamine—Forward: NCSI → NCNI → NRNI →IRNI→IRNL; Reverse IRNL→IRNI all steps in the preferred pathways—forward and reverse contain the S108N mutation. The mutation plays a central role in dictating the direction of evolution in both high drug (10^6^ *μM* pyrimethamine) and the drugless environment.

## IV. DISCUSSION

In this study, I introduce an abstraction called the mutation effect reaction norm (Mu-RN), that depicts how mutation effects and epistasis vary across environmental contexts. To demonstrate how it works, I apply previously developed mathematical methods introduced to measure higher-order epistasis on combinatorially complete data sets, across drug type and concentration. One of this study’s key messages are about how the capriciousness of mutation effects and interactions, as a function of environments and contexts, contributes to the complexity of the relationship between genotype and phenotype.

We should note the parallels between this perspective and classical debates about the “gene’s-eye view of evolution” between Sewell Wright and Ronald Fisher [3]. Though the specifics of Wright’s arguments were different than the ones outlined in this manuscript, he was an advocate of a more complex view of genetic systems, and critical of a simple, genic view of evolution [3, 4, 5]. The mutation effect reaction norm emerges from this intellectual tradition: it embodies complexity and the many types of interactions that define the shape of genetic systems and adaptive evolution.

### A. Tracking mutation effects across environmental contexts

With the mutation effect reaction norm, several questions can be examined, such as how multi-dimensional environments tune the phenotypic effects of mutations. We observe this through comparing the shape of the mutation effect reaction norms for a suite of mutations associated with resistance to pyrimethamine and cycloguanil (antifolate drugs used to treat malaria that are similar in structure) across a range of drug concentrations. In this case, we see how and why the adaptive landscapes differ for the two drugs: interactions between mutations in the *P. falciparum* dihydrofolate reductase protein backbone differ as a function of both drug type (pyrimethamine or cycloguanil) and drug concentration. Though pyrimethamine or cycloguanil have a similar mechanism of action and are similar in size (pyrimethamine molecular weight = 248.7; cycloguanil molecular weight = 251.7), these ostensibly subtle differences have meaningful consequences for patterns of resistance in nature [35].

Moving past the specific case of antimicrobial resistance, these findings speak to concepts that are of central importance in evolutionary theory. For example, the mutation effect reaction norm may inform models of how adaptive evolution occurs in fluctuating environments, where context-specific interactions can create opportunity and constraint [18, 27, 33]. These ideas are especially germane to modern efforts to improve on notions of static adaptive or fitness landscapes, towards the more realistic analogy of the fitness seascape [50].

### B. The Mu-RN and the dynamics of adaptation and reversal

The study’s detailed examination of one mutation’s interaction (S108N) serves as an example of how tracking the effects and interactions across environments allows one to identify (i) the specific contexts in which a given mutation is compensatory and (ii) the degree to which the effect is compensatory or not. The story that I have revealed about S108N likely applies to many “compensatory ratchet” mutations: their effects are often not binary—as in, they are beneficial in an environment or not—but rather, have stories that are more nuanced. Other examples include studies of bacterial translation machinery, where the contingent nature of compensatory mutations in evolution also manifests [51].

The findings surrounding “compensatory ratchet” mutations also highlights why reversal can be unlikely in circumstances where phenotypic effects of mutations (and interactions) are specific to certain environmental contexts. This has practical relevance and informs public health and biomedical approaches to addressing the antimicrobial resistance problem (for example). Studies have revealed why resistance management approaches that attempt to drive populations of resistant microbes “back” to more susceptible forms can be challenging [52, 53, 54]. And other studies have proposed methods and perspectives for how to reverse the effects of resistance [55]. We suggest that tracking mutation effects across environments, using the Mu-RN, can elucidate the molecular causes of irreversibility.

## V. CONCLUSION

The scientific world is full of models and abstractions that vary in their ability to describe relevant phenomena. What does the mutation effect reaction norm add? Is it just another abstraction that is engineered for clarifying purposes, but accomplishes the opposite? While only time will tell whether this new abstraction is useful, I have argued that it fills a notable gap, provides a simple, tractable, transposable depiction of how mutation effects and epistatic interactions can change across environmental contexts. This is consistent with prior descriptions of “environmental epistasis” [27], but made more generalizable, for potential applications across settings.

In order to depict environmental epistasis in simple terms, I have used the reaction norm as a basis for conception and comparison. Table 1 describes the differences between the reaction norm and the mutation effect reaction norm, including how one can interpret the information contained within them. The mutation effect reaction norm abstraction offers a different view of gene by environment interactions, by deconstructing genotype-phenotype maps in terms of the often peculiar interactions between genes and mutations across environments. This can offer a “mutation-centric” or “interaction-centric” view of genotype-phenotype maps where the object of interest is not entire haplotypes, but rather, the individual interactions between the mutations that compose those haplotypes, and environments. While evolutionary geneticists have long appreciated the importance of individual mutations, I argue that the mutation effect reaction norm appreciates another level of nuance, whereby we can author more detailed biographies of mutations and their interactions.

**Table 1:** Reaction norm vs. Mutation effect reaction norm. This table briefly describes the differences between the two abstractions

| Term | Data | Interpretation |
| --- | --- | --- |
| *Reaction norm* or norm of reaction | The performance, phenotype or trait value for different alleles, strains, mutants, variants, or forms, or populations, across an environmental gradient (continuous or other) | Used to identify and measure the effects of genes, environments, gene x environment interactions, phenotypic plasticity, and other related properties in quantitative genetics, evolutionary genetics, and ecology. |
| *Mutation effect reaction norm* (Mu-RN) | The phenotypic effect of individual or collections of mutations across a set of environments (continuous or other) | Used to measure how environments influence the effect of individual mutations, the interaction between suites of mutations, and epistatic interactions. It can help to explain how the environmental sculpts the topography of adaptive landscapes via environment-dependent mutation effects. |

Finally, I argue that the mutation effect reaction norm has important practical implications across several domains. By understanding how mutation effects and epistasis are driven by context, we can appreciate their role in obfuscating results from experimental studies of mutations [56], or in terms of how the context specificity of mutations may complicate genetic modification efforts [57]. For example, we should be careful to consider the environments in which the effects of engineered mutations evaluated.

The implications for predictive evolution are fairly obvious: the environmental mediator of nonlinear genetic interactions frames the topography of genotype-phenotype space, and by extension, how we expect the process of adaptive evolution to occur [58, 59, 60]. In addition, recent studies that support the role of contingency in evolution might be explained by the specificity of mutation effects, adaptive landscape topography or environmental context [61, 62]. In the biomedical arena, the abstraction has obvious connections to modern efforts to explain, predict or steer the evolution of antibiotic resistance, all of which involve an understanding of the effects of mutations[63, 64, 65, 66, 67].

Beyond drug resistance, the mutation effect reaction norm has practical applications to a range of other problems in biomedicine and public health. For example, the Mu-RN may allow us to better identify mutations in pathogens that are associated with emergence such as the ones identified in the form of a “watchlist” of mutations [68]. This notion has become especially relevant in the context of the COVID-19 pandemic. A year after the start of the pandemic, several variants of concern (VoC) began to circulate and define a new wave of the pandemic globally. These VoCs are the product of suites of mutations, some of which interact in a nonlinear fashion and have complicated our attempts to resolve which mutations are sole signatures of pathogenic potential (specifically, increased transmission and/or possible escape from vaccine-induced immunity) [69, 70]. I argue that the key to properly characterizing these mutations resides in an understanding of how environmental context shapes their effect. That is, host structure, demographics (e.g., age), and other factors may influence how a given SARS-CoV-2 mutation interacts with others, creating a variant of concern.

Notably absent from my introduction of the mutation effect reaction norm are analytical descriptions in the formal parlance of quantitative genetics. This is unlike the reaction norm, which has been the subject of these efforts in the past [71, 23, 72]. And the existence of analytically inspired studies in related topics, such as the use of rank orders of genotypes to infer genetic interactions [48, 39], or those that have examined the fate of mutations in fluctuating environments [73] suggest that similar formalisms may exist for the mutation effect reaction norm. This constitutes a future direction of investigation.

These gaps notwithstanding, the mutation effect reaction norm may encourage evolutionary geneticists to add nuance to conversations about how genotype relates to phenotype. Unidimensional questions about the contributions of a mutation to a phenotype are mostly insufficient. Moving forward, we should consider mutation effects with respect to the multidimensional environments which define the natural world.

## ACKNOWLEDGEMENTS

I would like to thank R. Oomen, M. Miller-Dickson, R. Guerrero, D. Weinreich, and P. Pennings for helpful discussions on this topic. I would like to acknowledge the editorial staff for helpful comments on a prior draft. I would like to thank M. Rebolleda-Gomez, R. Shaw, the participants, and other contributors to the symposium entitled “Society for the Study of Evolution (SSE) at 75 years: continuity and change in evolutionary research” at the 2021 Evolution Meetings. I would like to acknowledge speaker invitations from the following entities, where iterations of the ideas in this manuscript were discussed: University of Michigan; Princeton University; University of Connecticut; North Carolina State University; University of California, Santa Cruz; the University of Chicago; Lawrence University; the 2020 Population, Evolutionary, and Quantitative Genetics (PEQG) symposium (Genetics Society of America). I would like to acknowledge NSF Grant no. 1736253 “Using Biophysical Protein Models to Map Genetic Variation to Phenotypes.”

## SUPPORTING INFORMATION AND DATA ARCHIVING

Data and other materials can be found on Figshare: https://doi.org/10.6084/m9.figshare.16661371

## AUTHOR CONTRIBUTIONS

C.B.O. conceived the idea for the manuscript; curated, analyzed and interpreted the data; wrote and edited the manuscript.

## Notes

https://doi.org/10.6084/m9.figshare.16661371

### Competing Interest Statement

The authors have declared no competing interest.

### Summary of Updates

Slight alterations in phrasing and organization, more detailed methods. Color scheme revised to accommodate red-green color blindness.

## REFERENCES

[1] N.H. Barton, A.M. Etheridge, and A. Véber. The infinitesimal model: Definition, derivation, and implications. Theoretical Population Biology, 118:50–73, 2017.

[2] Evan A. Boyle, Yang I. Li, and Jonathan K. Pritchard. An expanded view of complex traits: From polygenic to omnigenic. Cell, 169(7):1177–1186, 2017.

[3] J Arvid Ågren. Sewall wright’s criticism of the gene’seye view of evolution. Evolution, 75(10):2326–2334, 2021.

[4] William B Provine. Sewall Wright and evolutionary biology. University of Chicago Press, 1989.

[5] Michael J Wade. Sewall wright: gene interaction and the shifting balance theory. Oxford surveys in evolutionary biology, 8:35–35, 1992.

[6] Timothy B. Sackton and Daniel L. Hartl. Genotypic context and epistasis in individuals and populations. Cell, 166(2):279–287, 2016.

[7] Daniel M Weinreich, Yinghong Lan, C Scott Wylie, and Robert B. Heckendorn. Should evolutionary geneticists worry about higher-order epistasis? Current Opinion in Genetics & Development, 23(6):700–707, 2013.

[8] Jeremy A. Draghi and Joshua B. Plotkin. Selection biases the prevalence and type of epistasis along adaptive trajectories. Evolution, 67(11):3120–3131, 2013.

[9] Hsin-Hung Chou, Nigel F. Delaney, Jeremy A. Draghi, and Christopher J. Marx. Mapping the fitness landscape of gene expression uncovers the cause of antagonism and sign epistasis between adaptive mutations. PLoS Genetics, 10(2):e1004149, 2014.

[10] Shimon Bershtein, Michal Segal, Roy Bekerman, Nobuhiko Tokuriki, and Dan S. Tawfik. Robustness–epistasis link shapes the fitness landscape of a randomly drifting protein. Nature, 444(7121):929–932, 2006.

[11] Tyler N. Starr and Joseph W. Thornton. Epistasis in protein evolution. Protein Science, 25(7):1204–1218, 2016.

[12] Amit Kumar, Chandrasekhar Natarajan, Hideaki Moriyama, Christopher C. Witt, Roy E. Weber, Angela Fago, and Jay F. Storz. Stability-mediated epistasis restricts accessible mutational pathways in the functional evolution of avian hemoglobin. Molecular Biology and Evolution, 34(5):1240–1251, 2017.

[13] Santiago F. Elena and Richard E. Lenski. Epistasis between new mutations and genetic background and a test of genetic canalization. Evolution, 55(9):1746–1752, 2001.

[14] R. Sanjuan and S. F. Elena. Epistasis correlates to genomic complexity. Proceedings of the National Academy of Sciences, 103(39):14402–14405, 2006.

[15] JM Cheverud and EJ Routman. Epistasis and its contribution to genetic variance components. Genetics, 139(3):1455–1461, 1995.

[16] H. J. Cordell. Epistasis: what it means, what it doesn’t mean, and statistical methods to detect it in humans. Human Molecular Genetics, 11(20):2463–2468, 2002.

[17] Xionglei He. The biology complicated by genetic analysis. Molecular Biology and Evolution, 33(9):2177–2181, 2016.

[18] Joanna Masel. Cryptic genetic variation is enriched for potential adaptations. Genetics, 172(3):1985–1991, 2006.

[19] David Haig. Does heritability hide in epistasis between linked SNPs? European Journal of Human Genetics, 19(2):123–123, 2010.

[20] O. Zuk, E. Hechter, S. R. Sunyaev, and E. S. Lander. The mystery of missing heritability: Genetic interactions create phantom heritability. Proceedings of the National Academy of Sciences, 109(4):1193–1198, 2012.

[21] Gibran Hemani, Sara Knott, and Chris Haley. An evolutionary perspective on epistasis and the missing heritability. PLoS Genetics, 9(2):e1003295, 2013.

[22] Rebekah A Oomen and Jeffrey Alexander Hutchings. Evolution of Reaction Norms. Oxford University Press, Oxford, England, 2020.

[23] Richard Gomulkiewicz and Mark Kirkpatrick. Quantitative genetics and the evolution of reaction norms. Evolution, 46(2):390, 1992.

[24] Rebekah A. Oomen and Jeffrey A. Hutchings. Genetic variability in reaction norms in fishes. Environmental Reviews, 23(3):353–366, 2015.

[25] Carl Schlichting and Massimo Pigliucci. Phenotypic Evolution: A Reaction Norm Perspective. Sinauer, Cary, NC, 1998.

[26] Carl D. Schlichting and Harry Smith. Phenotypic plasticity: linking molecular mechanisms with evolutionary outcomes. Evolutionary Ecology, 16(3):189–211, 2002.

[27] Haley A Lindsey, Jenna Gallie, Susan Taylor, and Benjamin Kerr. Evolutionary rescue from extinction is contingent on a lower rate of environmental change. Nature, 494(7438):463–467, 2013.

[28] Kenneth M. Flynn, Tim F. Cooper, Francisco B-G. Moore, and Vaughn S. Cooper. The environment affects epistatic interactions to alter the topology of an empirical fitness landscape. PLoS Genetics, 9(4):e1003426, 2013.

[29] Anne E. Hall, Kedar Karkare, Vaughn S. Cooper, Claudia Bank, Tim F. Cooper, and Francisco B.-G. Moore. Environment changes epistasis to alter trade-offs along alternative evolutionary paths. Evolution, 73(10):2094–2105, 2019.

[30] Elena R Lozovsky, Rachel F Daniels, Gavin D Heffernan, David P Jacobus, and Daniel L Hartl. Relevance of higher-order epistasis in drug resistance. Molecular Biology and Evolution, 38(1):142–151, 2020.

[31] Lingchong You and John Yin. Dependence of epistasis on environment and mutation severity as revealed byin SilicoMutagenesis of phage t7. Genetics, 160(4):1273–1281, 2002.

[32] Chuan Li and Jianzhi Zhang. Multi-environment fitness landscapes of a tRNA gene. Nature Ecology & Evolution, 2(6):1025–1032, 2018.

[33] Susanna K Remold and Richard E Lenski. Pervasive joint influence of epistasis and plasticity on mutational effects in escherichia coli. Nature genetics, 36(4):423–426, 2004.

[34] C. Brandon Ogbunugafor, C. Scott Wylie, Ibrahim Diakite, Daniel M. Weinreich, and Daniel L. Hartl. Adaptive landscape by environment interactions dictate evolutionary dynamics in models of drug resistance. PLOS Computational Biology, 12(1):e1004710, 2016.

[35] D. S. Peterson, W. K. Milhous, and T. E. Wellems. Molecular basis of differential resistance to cycloguanil and pyrimethamine in plasmodium falciparum malaria. Proceedings of the National Academy of Sciences, 87(8):3018–3022, April 1990.

[36] C. Brandon Ogbunugafor and Margaret J. Eppstein. Competition along trajectories governs adaptation rates towards antimicrobial resistance. Nature Ecology & Evolution, 1(1):1–8, 2016.

[37] Zachary R Sailer and Michael J Harms. Detecting high-order epistasis in nonlinear genotype-phenotype maps. Genetics, 205(3):1079–1088, 2017.

[38] Zachary R. Sailer and Michael J. Harms. Molecular ensembles make evolution unpredictable. Proceedings of the National Academy of Sciences, 114(45):11938–11943, 2017.

[39] Kristina Crona. Rank orders and signed interactions in evolutionary biology. eLife, 9:e51004, 2020.

[40] J. Otwinowski and J. B. Plotkin. Inferring fitness landscapes by regression produces biased estimates of epistasis. Proceedings of the National Academy of Sciences, 111(22):E2301–E2309, 2014.

[41] Zachary R. Sailer and Michael J. Harms. Uninterpretable interactions: epistasis as uncertainty. bioRxiv, preprint:1–32, 2018.

[42] Lorin Crawford, Ping Zeng, Sayan Mukherjee, and Xiang Zhou. Detecting epistasis with the marginal epistasis test in genetic mapping studies of quantitative traits. PLOS Genetics, 13(7):e1006869, 2017.

[43] Daniel M. Weinreich, Yinghong Lan, Jacob Jaffe, and Robert B. Heckendorn. The influence of higher-order epistasis on biological fitness landscape topography. Journal of Statistical Physics, 172(1):208–225, 2018.

[44] Frank J. Poelwijk, Vinod Krishna, and Rama Ranganathan. The context-dependence of mutations: A linkage of formalisms. PLOS Computational Biology, 12(6):e1004771, 2016.

[45] João V. Rodrigues, Shimon Bershtein, Anna Li, Elena R. Lozovsky, Daniel L. Hartl, and Eugene I. Shakhnovich. Biophysical principles predict fitness landscapes of drug resistance. Proceedings of the National Academy of Sciences, 113(11):E1470–E1478, 2016.

[46] Rafael F Guerrero, Samuel V Scarpino, João V Rodrigues, Daniel L Hartl, and C Brandon Ogbunugafor. Proteostasis environment shapes higher-order epistasis operating on antibiotic resistance. Genetics, 212(2):565–575, 2019.

[47] Victor A. Meszaros, Miles D. Miller-Dickson, and C. Brandon Ogbunugafor. Lexical landscapes as large in silico data for examining advanced properties of fitness landscapes. PLOS ONE, 14(8):e0220891, 2019.

[48] Daniel M Weinreich. The rank ordering of genotypic fitness values predicts genetic constraint on natural selection on landscapes lacking sign epistasis. Genetics, 171(3):1397–1405, 2005.

[49] C. Brandon Ogbunugafor and Daniel Hartl. A pivot mutation impedes reverse evolution across an adaptive landscape for drug resistance in plasmodium vivax. Malaria Journal, 15(1):1–20, 2016.

[50] Ville Mustonen and Michael Lässig. From fitness landscapes to seascapes: non-equilibrium dynamics of selection and adaptation. Trends in Genetics, 25(3):111–119, 2009.

[51] Sandeep Venkataram, Ross Monasky, Shohreh H. Sikaroodi, Sergey Kryazhimskiy, and Betul Kacar. Evolutionary stalling and a limit on the power of natural selection to improve a cellular module. Proceedings of the National Academy of Sciences, 117(31):18582–18590, 2020.

[52] Maria Sjölund, Karin Wreiber, Dan I. Andersson, Martin J. Blaser, and Lars Engstrand. Long-term persistence of resistant enterococcus species after antibiotics to eradicate helicobacter pylori. Annals of Internal Medicine, 139(6):483–487, 2003.

[53] Dan I. Andersson and Diarmaid Hughes. Antibiotic resistance and its cost: is it possible to reverse resistance? Nature Reviews Microbiology, 8(4):260–271, 2010.

[54] M. Sundqvist, P. Geli, D. I. Andersson, M. Sjolund-Karlsson, A. Runehagen, H. Cars, K. Abelson-Storby, O. Cars, and G. Kahlmeter. Little evidence for reversibility of trimethoprim resistance after a drastic reduction in trimethoprim use. Journal of Antimicrobial Chemotherapy, 65(2):350–360, 2009.

[55] M. Baym, L. K. Stone, and R. Kishony. Multidrug evolutionary strategies to reverse antibiotic resistance. Science, 351(6268):aad3292–aad3292, 2015.

[56] Tom J. Little and Nick Colegrave. Caging and uncaging genetics. PLOS Biology, 14(7):e1002525, 2016.

[57] Peter B. Otoupal, Keesha E. Erickson, Antoni Escalas-Bordoy, and Anushree Chatterjee. CRISPR perturbation of gene expression alters bacterial fitness under stress and reveals underlying epistatic constraints. ACS Synthetic Biology, 6(1):94–107, 2016.

[58] Adam C. Palmer and Roy Kishony. Understanding, predicting and manipulating the genotypic evolution of antibiotic resistance. Nature Reviews Genetics, 14(4):243–248, 2013.

[59] J. Arjan G.M. de Visser and Joachim Krug. Empirical fitness landscapes and the predictability of evolution. Nature Reviews Genetics, 15(7):480–490, 2014.

[60] Allison J. Lopatkin and James J. Collins. Predictive biology: modelling, understanding and harnessing microbial complexity. Nature Reviews Microbiology, 18(9):507–520, 2020.

[61] Mateusz Kędzior, Amanda K. Garcia, Meng Li, Arnaud Taton, Zachary R. Adam, Jodi N. Young, and Betül Kaçar. Molecular foundations of precambrian uniformitarianism. bioRxiv, preprint:1–24, 2021.

[62] Victoria Cochran Xie, Jinyue Pu, Brian PH Metzger, Joseph W Thornton, and Bryan C Dickinson. Contingency and chance erase necessity in the experimental evolution of ancestral proteins. eLife, 10:e67336, 2021.

[63] Adam C. Palmer, Erdal Toprak, Michael Baym, Seungsoo Kim, Adrian Veres, Shimon Bershtein, and Roy Kishony. Delayed commitment to evolutionary fate in antibiotic resistance fitness landscapes. Nature Communications, 6(1):1–8, 2015.

[64] Jennifer Knies, Fei Cai, and Daniel M. Weinreich. Enzyme efficiency but not thermostability drives cefotaxime resistance evolution in TEM-1 β-lactamase. Molecular Biology and Evolution, 34(5):1040–1054, 2017.

[65] Erida Gjini and Kevin B Wood. Price equation captures the role of drug interactions and collateral effects in the evolution of multidrug resistance. eLife, 10:e64851, 2021.

[66] Sarah M. Ardell and Sergey Kryazhimskiy. The population genetics of collateral resistance and sensitivity. bioRxiv, preprint:1–39, 2020.

[67] Daniel Nichol, Peter Jeavons, Alexander G Fletcher, Robert A Bonomo, Philip K Maini, Jerome L Paul, Robert A Gatenby, Alexander RA Anderson, and Jacob G Scott. Steering evolution with sequential therapy to prevent the emergence of bacterial antibiotic resistance. PLoS computational biology, 11(9):e1004493, 2015.

[68] Craig R. Miller, Erin L. Johnson, Aran Z. Burke, Kyle P. Martin, Tanya A. Miura, Holly A. Wichman, Celeste J. Brown, and F. Marty Ytreberg. Initiating a watch list for ebola virus antibody escape mutations. PeerJ, 4:e1674, 2016.

[69] Sarah P. Otto, Troy Day, Julien Arino, Caroline Colijn, Jonathan Dushoff, Michael Li, Samir Mechai, Gary Van Domselaar, Jianhong Wu, David J.D. Earn, and Nicholas H. Ogden. The origins and potential future of SARS-CoV-2 variants of concern in the evolving COVID-19 pandemic. Current Biology, 31(14):R918–R929, 2021.

[70] Carolina Lucas, Chantal BF Vogels, Inci Yildirim, Jessica E Rothman, Peiwen Lu, Valter Monteiro, Jeff R Gehlhausen, Melissa Campbell, Julio Silva, Alexandra Tabachnikova, et al. Impact of circulating sars-cov-2 variants on mrna vaccine-induced immunity. Nature, pages 1–7, 2021.

[71] G. de Jong. Quantitative genetics of reaction norms. Journal of Evolutionary Biology, 3(5-6):447–468, 1990.

[72] Carl D. Schlichting and Massimo Pigliucci. Gene regulation, quantitative genetics and the evolution of reaction norms. Evolutionary Ecology, 9(2):154–168, 1995.

[73] Ivana Cvijović, Benjamin H. Good, Elizabeth R. Jerison, and Michael M. Desai. Fate of a mutation in a fluctuating environment. Proceedings of the National Academy of Sciences, 112(36):E5021–E5028, 2015.

